# Toggling *Geobacter sulfurreducens* metabolic state reveals hidden behaviour and expanded applicability to sustainable energy applications

**DOI:** 10.1101/623652

**Authors:** Mir Pouyan Zarabadi, Steve J. Charette, Jesse Greener

## Abstract

Electroactive biofilms are under intense scrutiny due to their potential to enable new sustainable technologies for energy production and bioremediation. However, severely reduced metabolic activity at low concentrations is a barrier to their implementation. A microfluidic approach was used for real-time respiration experiments on a *Geobacter sulfurreducens* biofilm to overcome these constraints. Precise changes to solution conditions enabled rapid and reversible switching between biofilm metabolic states, leading to the following discoveries. (i) Flow reactors can maintain biofilm activity at concentrations as low as 15 µM; (ii) a “pseudo-active” metabolic state separates active and inactive states; and (iii) acetate conversion can be as high as 90 percent for active biofilms at the pseudo-activity threshold.

---

Electroactive biofilms (EABs) have the potential to accelerate the development of new sustainable processes including the conversion of waste water to electricity.^1^ In addition EABs have been demonstrated for synthesis of chemical compounds and nanomaterials^2-4^ and environmental remediation.^5^ Moreover, new work is underway to understand and optimise their coupling with other microorganisms to expand functionality, via direct interspecies electron transfer.^5-7^ EABs from *Geobacter sulfurreducens* are among the most studied because after the oxidation of an acetate molecule, electrons are efficiently conducted to external electron acceptors, such as an electrode via a direct externally electron transfer mechanism.^6,9^ In studies using a microbial fuel cell (MFC) or a microbial electrolysis cell (MEC), the liquid phase typically contains high concentration of organic molecules, well in excess of 120 mg·L^-1^ (equivalent to 2 mM acetate). In contrast, far fewer studies have focused specifically on *Geobacter* performance at low concentrations, such as those occurring in selected natural environments where these organisms are found.^10-11^ Recently, the importance of the low-concentration regime has been highlighted by a practical limitation that strongly reduces EAB current in MFCs at chemical oxygen demand (COD) levels between 100 to 200 mg·L^-1^ and outright at 50 mg·L^-1^.^12,13^ The reason for this loss of activation is not known, but the impact is clear. In addition to poor performance at low concentrations, subsequent processing steps will be required to reduce final concentrations down to environmental discharge limits, adding another hurdle to implementation and increasing complexity.^14^

In the 1940’s, Monod and others studied the effect of external influences such as nutrient concentrations on bacterial growth and metabolic activity.^15^ Forty years later, detailed studies into nutrient mass transfer into biofilms and their effects on metabolism were conducted.^16^ However, the difficulty in obtaining sufficient control over reaction parameters continues to limit studies of biofilm metabolic activity at low concentrations. For example, in bulk electrochemical cells, solution concentration continuously changes as acetate is oxidised and by-products are produced. To avoid concentration cycling, flow systems can be used to provide a continuous nutrient supply. Hydrodynamic conditions are also known to impact molecular transport via their effect on diffusion barriers around the EAB^17^ and potentially from forced convection through voids, cavities and channel structures, as noted previously in non-electroactive biofilms.^18-23^ It also appears that molecular transport into the highly active biofilm regions near the electrode surface plays a significant role under low-concentration conditions.^24^ Taken together, well-controlled flow systems have a great potential for discovery and optimisation of EABs at low concentrations. A centimetre-scale flow system demonstrated the potential for studies of current generation at low concentrations.^25^ However, dead volumes would limit response time and its demonstrated control in the range of 0 to 2.3 mM acetate is needed for the present study. Millifluidic systems can be easily constructed using standard machining techniques or even with standard flexible tubing. For example, respirometric studies of *Pseudomonas* biofilms using gas-permeable silicon tubing and a CO_2_ detection system^26-31^ could in principle be adapted for electrode-adhered anaerobic EABs, but the integration of electrodes within the tubing interior and the requirement of an oxygen-purged environment would present difficult challenges. Millifluidic systems and larger must also force a balance between reasonable solution volumes and limitations to attainable flow velocities or applied shear stresses. Alternatively, microfluidic approaches combine the advantages of low material consumption over a large range of applied hydrodynamic conditions, established methods for electrode integration and high control of chemical concentrations via on-chip dilution,^32-33^ microvalves,^34^ and formation of concentration gradients.^35^ Increasingly, microfluidic flow cells are recognised as viable platforms for precision bioanalytical chemistry-based studies of biofilms, including hydrodynamic voltammetry.^36^ For example, our group recently developed a robust three-electrode microfluidic flow cell for electrochemical measurements of bacterial biofilms from *Pseudomonas*^35^ and *Geobacter* species.^38^ Such a platform combines the benefits of direct and continuous measurements of bacterial metabolism^36^ with standard microfluidic advantages such as unsurpassed control over solution-phase hydrodynamic and chemical properties, as well as low liquid consumption.^39^ In addition, complex multi-step experimental sequences can be devised with vanishingly low latency, thus enabling more precise and rapid studies of microbiological systems than ever before.^40-42^

In this work, we used a microfluidic three-electrode electrochemical flow cell to measure the current from *G.sulfurreducens* EABs via chronoamperometry, while low acetate concentrations were applied at well-defined flow rates. Using this setup, we identified threshold conditions that separated active, pseudo-active and inactive metabolic states. Moreover, these thresholds were dependent on flow rate. Figure 1a shows the schematic of the microfluidic system. The PCA strain of *G. sulfurreducens* was inoculated into the microchannel for 3 h with a 0.5 mL·h^-1^ flow rate. A three-way liquid mixer was placed between the microchannel inlet and three different air-tight glass syringes containing acetate concentrations of [Ac]_1_ = 10, [Ac]_2_ = 0.2 and [Ac]_3_ = 0 mM. With the use of one-way stopcock valves at the inlet side of mixer inlets 1-3 (MI_1_, MI_2_, MI_3_) and flow control via the syringe pumps, the final nutrient concentration could be determined from real-time dilutions in the range of 10 to 0 mM based on the formula:

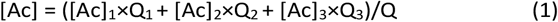

**Fig. 1.**
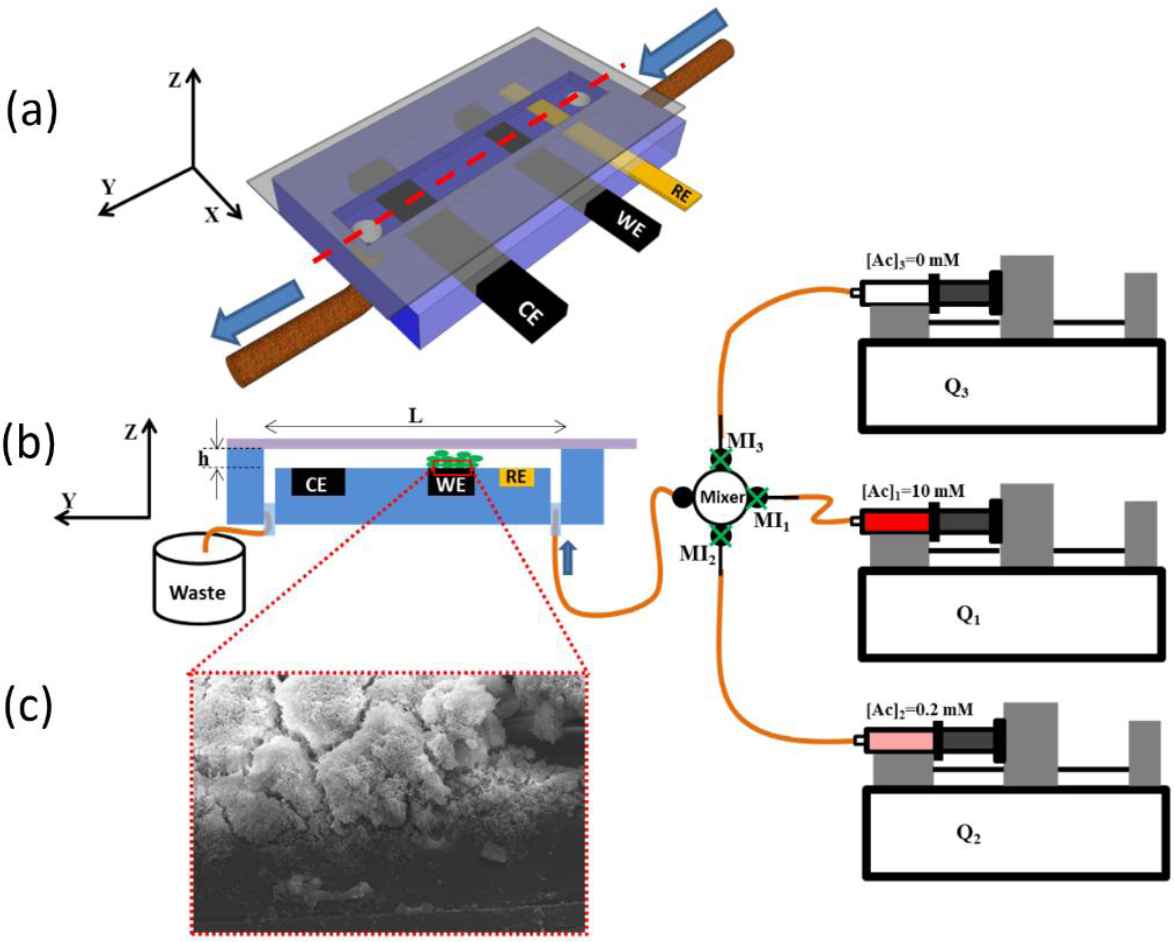
Experimental setup, (a) Three-dimensional schematic of a microfluidic electrochemical flow cell showing the gold reference electrode (RE), graphite working electrode (WE), and counter-electrode (CE). The height, width and length are h = 400 μm, w = 2000 μm and L = 3 cm, respectively. All electrode portions exposed to the channel have the same width as the channel and extend for 3000 μm along the channel length. The dashed line shows the position of the cross-section for (b). (b) 2D schematic of the device cross-section (y-z plane) with three syringe pumps holding 50 mL gas-tight syringes containing nutrient solutions of 0, 0.2 and 10 mM acetate. The outlet of each syringe is connected to one inlet of a mixing element via a one-way stopcock valve. The mixer outlet is connected to the device inlet. The inoculum syringe is not shown, (c) SEM image of *G. sulfurreducens* biofilm (>1000 h age) on the graphite working electrode following the experiments described in this work. All experiments were conducted at 22 °C.

where the total flow rate is given by Q = Q_1_ + Q_2_ + Q_3_. Thus the average liquid velocity within the channel could be calculated from *v* = Q/A, where A is the channel cross-sectional area based on the channel height (h = 400 µm) and width (w = 2 mm). In this manner, rapid and precise changes in both [Ac] and Q could be generated on demand within a matter of seconds and maintained for durations ranging from seconds to tens of hours. After passing from the mixer to the microchannel, the liquid first flowed past a gold (pseudo) reference electrode (RE), followed by a graphite working electrode (WE), and finally a counter electrode (CE). The surface area of all electrodes exposed to the liquid solution was 3 mm × 2 mm.

Additional information on design and fabrication of the three-electrode microfluidic device can be found in our previous work^37,38^ and in the Supporting Information. Because the electroactive *G. sulfurreducens* bacteria must externally transfer electrons to complete the tricarboxylic acid metabolic cycle,^38,43^ their growth was strictly limited to the WE (Figure 1), where biofilm growth and acetate turnover rate was monitored directly via chronoamperometry. Placement of the RE in the upstream position ensured that by-products could not diffuse upstream with speed sufficient to overcome the downstream flow velocity (Supporting Information, Section 2). Thus, contamination of the RE environment was stable for the duration of the experiment, as demonstrated previously.^36-38^ The electric current is related to the overall biofilm metabolism based on the fact that oxidation of one mole of acetate (mol_Ac_) produces 8 moles of free electrons (mol_e_).^38,44^ Therefore, the current, I (C·s^-1^), can be converted to the acetate turnover rate, 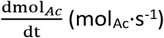, based on Equation 2:

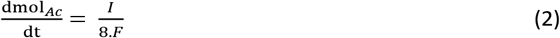

where F is Faraday’s constant, F=9.6485 × 10^4^ C·mol_e_^-1^ and 8 is the proportionality constant for moles of electrons produced to moles of acetate oxidised. The combination of real-time control over [Ac] and Q together with instantaneous measurement of the acetate turnover rate presents a simple but powerful approach to study changes in EAB metabolism. In the remainder of this paper, we quantified the EAB metabolic activity via electric current.^45^

As shown in the Supporting Information (Figure S2a), after channel inoculation and growth for approximately 300 h at [Ac] = 10 mM, the current reached 37.5 ± 0.75 µA (6.16 ± 0.12 A·m^-2^). After this time, the current produced by the biofilm supplied with a nutrient medium of [Ac] = 10 mM at Q = 0.25 mL·h^-1^ remained the same until the very end of the experiment, immediately before decommission of the flow cell. Therefore, we deduced that after 300 h the biofilm was mature and its thickness was constant.^46-48^ SEM measurements of the biofilm at the end of the experiment showed that the biofilm had an average height of 80 µm. This resulted in a constriction in the flow channel, resulting in an average headspace above the biofilm of 320 µm. After verifying that the current was stable for another 210 h, the experiments were started.

To test the response of a fully mature biofilm to different degrees of nutrient limitation, we switched the system to low-concentration mode by closing valve MI_1_ and opening valves MI_2_ and MI_3_. In this step, we varied [Ac] between 0 and 0.2 mM (200 µM) while maintaining Q = 0.25 mL·h^-1^ based on a simplified form of Eqn. (1): [Ac] = ([Ac]_2_×Q_2_ + [Ac]_3_×Q_3_)/Q. The upper limit of 200 µM was chosen based on the lowest concentration which could maintain full EAB activity at the lowest flow rate used in this study (0.25 mL·h^-1^). Switching to [Ac] = 200 µM resulted in an immediate reduction in current, which continued for nearly 1 h until a steady-state value of approximately 10 µA (1.67 A·m^-2^) was reached (Supporting Information, Figure S2b). Subsequently, we reduced [Ac] in a stepwise manner from 200 to 140 µM and eventually to 80 µM. As shown in Figure 2a, dropping the concentration to [Ac] = 140 µM resulted in a steady reduction in current from region I levels to approximately 4 µA within the first few minutes. Unlike in other studies where current decreased monotonically to zero after applying zero concentration nutrient solutions,^38,49^ or where a new lag phase is induced after a switching acetate for a complex carbon source,^50^ the switch to the low but non-zero acetate concentrations used here, induced a fluctuating current (region II). The fluctuating state could be maintained for up to 40 hours, as long as very low, but non-zero concentrations were applied (Supporting Information, Figure S5c). Based on the power spectrum from a fast Fourier transform analysis of the time series data, the frequency component contributing to these oscillations could be as high as 2 h^-1^. This is equivalent to periodicity of 30 mins. Thus, according to the Nyquist sampling theorem, the wave form was properly represented by the sampling rate which was 1/30 s (Supporting Information, Section 1). In contrast, the current profiles recorded during operation at nutrient concentrations at or above [Ac] = 200 µM featured nearly unobservable fluctuations in EAB current intensity and high frequency components that were 40 times slower than those in region II. Supporting Information, Section 7 contains more information on the stark differences in current profiles between regions I and II. Switching to [Ac] = 80 µM while maintaining Q= 0.25 mL·h^-1^ resulted in a current reduction to a stable, non-fluctuating value of nearly 1 µA (region III), which matched the background current measured separately for the same setup. Further reduction of [Ac] had no effect on electrical current, indicating that the background current had indeed been achieved in the last step. Therefore, three distinct metabolic activity states could be selected based on [Ac] and from the EAB current profiles: an active state (I), a pseudo-active state (II), and an inactive state (III) (Figure 2a).

**Fig. 2.**
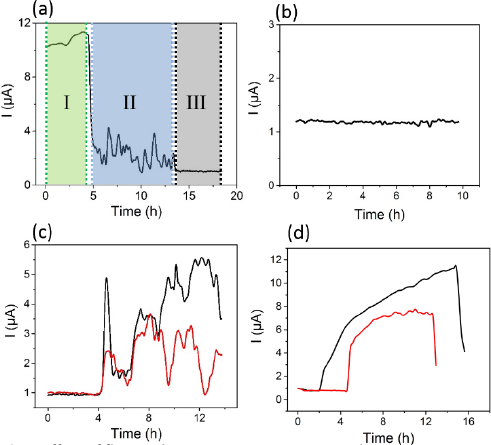
Effect of flow and nutrient concentration on the current output of a fully mature *G. sulfurreducens* biofilm, (a) Current profiles due to changes in [Ac]. Region I shows an active biofilm exposed to [Ac] = 200 μM (green). Region II shows a pseudo-active biofilm with exposure to [Ac] = 140 μM acetate (blue). Region (III) shows an inactive biofilm after exposure to [Ac] =80 μM (grey). In all cases, Q = 0.25 mL·h^-1^. (b) Constant background current levels recorded during increases to [Ac] in 10 μM increments while [Ac]< [AC]_PA_ with Q = 0.25 mL·h^-1^. (c) Current output following two separate transitions from inactivity (at [Ac] = 0) to pseudo-activity via [AC]_PA_ = 160 μM with Q = 0.25 mL·h^-1^ (black curve) and [AC]_PA_ = 30 μM acetate at Q = 1 mL·h^-1^ (red curve), (d) Current output following two separate transitions from inactivity to full activity via [AC]_A_ = 200 μM with Q = 0.25 mL·h^1^ flow rate (black line) and [AC]A = 40 nM acetate with Q = 1 mL·h^1^ flow rate (red line). The reduction in current observed in both data sets was the result of a switch to [Ac] = 0 M.

Taking advantage of the ability to impose rapid, accurate changes in experimental conditions, an iterative screening procedure was developed to determine the exact transition concentrations that separated activity states I, II and III. This was accomplished via the application different [Ac] pulses while maintaining a constant Q. Specifically, in each iteration, concentration was cycled between [Ac] = 0 and an elevated concentration [Ac]_i_ (where the subscript i is an index value for each iteration) until the EAB current increased above the background. The increase in concentration from one pulse to the next was [Ac]_i+1_ -[Ac]_i_ = 10 µM. At concentrations too low to provoke a change from the metabolically inactive state, no deviation from the background current was observed (Figure 2b). In separate experiments, it was verified that the inactive state could be maintained for at least 20 h at such low, but non-zero concentrations (data not shown). With continued stepwise increases in [Ac]_i_, a threshold concentration [Ac]_PA_ was identified that resulted in the transition to the pseudo-active state. This transition was marked by an increase in current from background levels to the same fluctuating state observed in region II of Figure 2a. Continued increases to [Ac]_i_ during subsequent iterations resulted in similar transitions to pseudo-activity until eventually, a new transition behaviour occurred in which the increase in current from background was smooth and asymptotically approached a maximum current I_max_. At and above this concentration, the biofilm could achieve continuous metabolic activity, thus defining a second concentration threshold for fully active EABs, [Ac]_A_.

As flow is known to affect the performance of *G. Sulfurreducens*,^38,51^ measurements of [Ac]_PA_ and [Ac]_A_ were repeated in triplicate at 5 different total flow rates in the range of Q = 0.25 to 1.5 mL·h^-1^. This corresponded to flow velocities of approximately v = 5 to v = 30 mm·min^-1^ and τ = 1.2 to 7 mPa (see Supporting Information, Section S2, Table S1 for more information). Figures 2c and 2d show typical transitions following a switch from [Ac] = 0 to [Ac]_PA_ and to [Ac]_A_, respectively. In all cases, the threshold concentrations decreased as Q increased. Due to the unsteady nature of the pseudo-active metabolic state, we could not confirm whether the flow rate affected the current output profile after transitions from [Ac] = 0 µM to [Ac]_PA_ (Figure 2c). However, as observed in Figure 2d, transitions from [Ac] = 0 to [Ac]_A_ at different values of Q did showed differences in the current rise time and I_max_. Specifically, for low [Ac]_A_ at high Q, the rise time was short and I_max_ was low, whereas for higher [Ac]_A_ at low Q, the rise time was slow, but I_max_ was larger. Increasing Q at any given [Ac]_A_ resulted in shorter rise times and higher I_max_. Thus Q appeared to control the rise time, whereas [Ac] and Q both affected I_max_.

Figure 3 summarises the results from the experiments described above. In this case, three distinct biofilm metabolic activity states were separated by threshold concentrations [Ac]_PA_ and [Ac]_A_. Both threshold concentrations were reduced with increasing Q, with good repeatability in their measured values. To the best of our knowledge, the pseudo-active metabolic state separating the active and inactive states has never been observed before. Another major observation is that increasing Q can maintain metabolic activity at much lower [Ac] than in bulk conditions, paving the way for exploration of simple flow-based techniques to reduce the previously observed lower concentration limits. For example, the previously reported loss of EAB activity at nutrient concentrations of approximately 50 mg·L^-1^ COD (equivalent to 0.83 mM acetate)^13^ is 30 and 55 higher than measured thresholds [Ac]_A_ and [Ac]_PA_ at high flow rate. It is noted that an exponential extrapolation of our results to static conditions (Q = 0 mL·h^-1^) does predict [Ac]_PA_ near 0.8 mM, confirming the pertinence of this work to previous studies. We also noted that the present measurements were conducted via chronoamperometry, which applies a constant potential throughout the experiment. This rules out a reduction in electrostatic-driven diffusion as the mechanism responsible for the loss of biofilm activity, which was proposed as a possible explanation for threshold concentrations in bulk microbial fuel cells.^13^

**Fig. 3.**
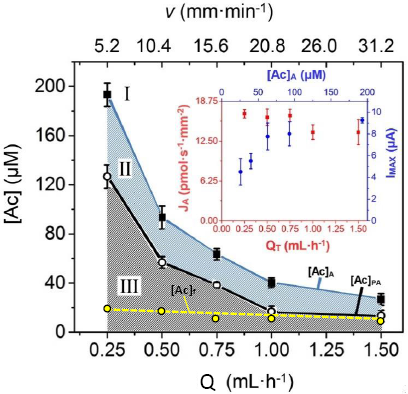
Dependency of concentration thresholds [AC]_PA_ (hollow circles) and [AC]_A_ (solid squares) on Q and equivalent flow velocity (V_A_) for a fully mature biofilm of *G. sulfurreducens* bacteria. The upper portion of the plot labelled I (white) indicates [Ac] and Q conditions resulting in active biofilm metabolism. The intermediate band labelled II (blue crosshatch) indicates [Ac] and Q conditions resulting in pseudo-active biofilm metabolism. The lower region of the plot labelled III (black crosshatch) indicates [Ac] and Q conditions resulting in biofilm inactivity. The yellow circles and respective trend line (dashed) are the calculated [Ac]_f_ following acetate consumption at the WE. Error bars were not added because they were smaller than the data points. Inset: Nutrient flux threshold J_A_ (pmol·s^-1^) as a function of Q (red) and maximum current Output I_max_ (μA) as a function of [AC]_A_ (blue). Error bars for all points were generated from three separate experiments at the same flow rate.

To better understand the role of flow in determining the loss of full metabolic activity, we converted the threshold concentration [Ac]_A_ and the corresponding flow conditions to the threshold convective acetate flux through the channel:

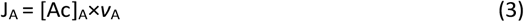

where *v*_A_ is the corresponding linear flow velocity to a specific [Ac]_A_ value. As shown in the inset plot in Figure 3, J_A_ is nearly constant at 15.6 pmol·s^-1^·mm^-2^ for all flow velocities. Therefore, it is concluded that J_A_ is the influential parameter in determining the metabolic activity state and not independent values of [Ac] or *v*. We note that the range in the calculated velocity values in Figure 3 apply to an empty channel. However, both EAB and non-electroactive biofilm thicknesses are known to increase under imposed fluid flow shear, which could affect local flow velocity.^52,53^ According to the SEM results discussed earlier, the reduced head space above the EAB, resulted in local velocities impinging on the upstream side of the biofilm being 18.7 pmol·s^-1^·mm^-2^. Unlike J_A_, it was noted that I_max_ was not constant over all *v*_A_ values. As observed in the Figure 3 inset, I_max_ increased with increasing [Ac]_A_ (and lower *v*_A_), similar to the observation related to Figure 2d. This is consistent with the expectation that solution-phase nutrient molecules should diffuse more rapidly into the EAB at high [Ac] because of higher concentration gradients between the bulk liquid and the depletion layer near the EAB surface.

The final acetate concentration, [Ac]_f_, after catalytic oxidation by EABs is an important performance measure for MFCs and other bioelectrochemical systems for environmental remediation. Following liquid contact with the EAB, [Ac]_f_ was determined for different initial acetate concentrations, [Ac]_0_, and flow rates using Equation 4. Acetate conversion efficiency, ε_Ac_, was determined by 5.

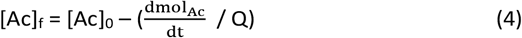

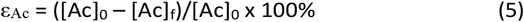

Here we evaluate [Ac]_f_ and ε_Ac_ for different [Ac]_A_ values shown in Figure 3. That is, we used Eqs. 4 and 5, with [Ac]_0_ = [Ac]_A_, I_max_ values obtained from data presented in Figure 3 (inset). Interestingly, the results for [Ac]_f_ are plotted in Figure 3 were approximately 10-20 µM for all applied threshold concentrations, [Ac]_A_. These corresponded to ε_Ac_ in the range of 60% (at high Q) to 90% (at low Q). We do not believe that relatively low ε_Ac_ at high Q is related to reduced contact time between the biofilm and the flowing liquid, as this would not explain why [Ac]_f_ is nearly constant with Q, even slightly decreasing as Q increased. Instead, it appears that these final concentrations are approaching a lower limit, which may be either fundamental to the biological system or related to our setup. As a comparison, we conducted experiments at higher concentrations ranging from [Ac]_0_ = 0.3 to [Ac]_0_ = 10 mM in the same flow rate range used previously. In this case, the calculated ε_Ac_ values were much lower (Supporting Information). Using average EAB currents achieved during [Ac]_0_ = [Ac]_PA_, ε_Ac_ was also determined to also be slightly worse than for [Ac]_0_ = [Ac]_A_, though still much better than for [Ac] > [Ac]_A_. Therefore, it is concluded that the best conversion was achieved for [Ac]_A_. See Supporting Information (Section 8, Figure S5) for further discussion on this point, including an evaluation of the role of liquid/EAB contact time on ε_Ac_. It is worth noting that the resulting final concentrations are equivalent to 0.65-1.25 mg·L^-1^, which is well-below the previously mentioned regulatory limits of 30 mg·L^-1^. The [Ac]_f_ leaving the device were generally lower than the lower limit for biofilm pseudo-activity ([Ac]_PA_), except at the highest flow rates where the two concentration profiles began to converge. The apparent contradiction that [Ac]_f_ can be lower than [Ac]_PA_ should be considered in the context of the experiment. The threshold [Ac]_PA_ was the minimum concentration required to transition an inactive biofilm to pseudo-activity. In the current case, biofilms are supplied sufficient nutrients to establish some level of metabolic activity (i.e., [Ac]_0_ ≥ [Ac]_PA_). Therefore, the observation that [Ac]_f_ was generally lower than [Ac]_PA_ is interpreted to mean that the entire biofilm could maintain its activity state, even in the downstream positions where concentrations were reduced by continuous acetate oxidation during the liquid/biofilm contact time. This is an interesting point that should be investigated further.

The present work opens the door to fundamental questions related to low activity thresholds and excellent acetate conversion in microflow conditions. In addition to a reduction in the diffusion barriers in flowing systems, other factors might play a role. For example, new evidence of flow-based biofilm deacidification^38^ or the potential for convective flow through the biofilm to deliver nutrient molecules to deeper and more electrically active portions of the EAB may also play a role.^54^ The root cause leading to loss of full activity should be also be investigated, with a particular focus on the mechanism causing the curious current oscillations in the pseudo-active metabolic state. The common [Ac]_f_ for all [Ac]_A_ threshold values (and their corresponding flow rates) should also be investigated. Simple modifications to the setup such as highly parallel channels and electrodes with greater surface areas and longer liquid/biofilm contact times should be investigated as approaches to potentially further reduce [Ac]_f_ and improve current generation. In doing so, the true biological lower [Ac] limit to maintain activity may also be discovered. Finally, a microfluidic MFC should be tested at low-concentrations to determine the extent to which the excellent figures of merit for a microfluidic three-electrode device transfer to other bioelectrochemical systems.

## Conclusions

A microfluidic electrochemical flow cell was used to study the metabolic state of electrode-adhered *Geobacter sulfurreducens* electroactive biofilms via chronoamperometry. The real-time current response to changes in acetate concentration and solution flow rate enabled the identification of three separate metabolic states: an active state, an inactive state and a newly identified pseudo-active state. In addition, flow conditions could strongly suppress loss of metabolic activity at low concentrations. Specifically, full biofilm activity could be maintained at concentrations as low as [Ac] = 25 µM, while pseudo-activity was maintained at concentrations as low as [Ac] = 15 µM with flow rates of 1.5 mL·h^-1^ (ca. 30 mm·min^-1^). This represents more than 50 times reduction in lowest usable Ac concentrations compared to bulk MFC studies. The final acetate concentrations leaving the device could reach as low as 10 µM or equivalently 0.65 mg·L^-1^ COD. Therefore, by extending previous flow studies to microscale flows of low concentration acetate solutions, this work paves the way for new approaches to optimisation and study of bioelectrochemical systems involving *Geobacter* and other electroactive biofilms.

## Contributions

MPZ helped to design the experimental methodology, conducted all experiments, performed data analysis and assisted in preparation of the manuscript. SC provided expertise on microbiological protocols and assisted with the manuscript. JG helped to design the experimental methodology and data analysis and wrote the manuscript.

## Acknowledgements

The authors thank Luc Trudel and Laurent Smith (U. Laval) for their help with culture of *G. sulfurreducens* and Richard Janvier (U. Laval) for technical assistance with SEM. This research was supported by generous funds from the Natural Sciences and Engineering Research Council of Canada, the Canadian Foundation for Innovation and Sentinelle Nord. Greener is the recipient of an early researcher award from the Fonds de recherche du Québec -nature et technologies. Greener and Charette are supported by a joint FRQNT AUDACE grant (high risk, high reward) to study microbiological systems using microfluidics. The authors also thank Molly K. Gregas for assistance with technical edits.

