## Supplemental Information for "Toggling *Geobacter sulfurreducens* metabolic state reveals hidden behaviour and expanded applicability to sustainable energy applications"

### Sections:

1. Device fabrication and experiment methodology

2. Hydrodynamic considerations at and away from the working electrode

3. Bacterial preparation

4. Reference electrode calibration and stability

5. Device inoculation and growth

6. Identification of cytochrome c formal potential by cyclic voltammetry

7. Fluctuations in electrical current in different metabolic activity states

8. Efficiency of acetate digestion

9. References

### 1. Device fabrication and experiment methodology

In this work, a 3 electrode microfluidic flow cell was fabricated in a glass-sealed elastomeric chip which was housed in an anaerobic chamber. The working and counter electrodes were cut into 3 mm × 20 mm strips from a 10 cm x 10 cm graphite sheet. A gold pseudo reference electrode was made by electroless deposition on a polystyrene substrate and cut into 3 mm × 20 mm. A negative photoresist was used for device fabrication. The electrodes were placed on photoresist features and were covered with a mixture of liquid polydimethylsiloxane (PDMS) and cross-linking agent Sylgard184 (Dow Corning, Canada) in 10:1 ratio. After curing for 4 h at 70 °C, the device was taken from the mold and the electrodes and microchannel were cleaned and sterilized with 1M HCl and 70% EtOH, respectively. The inlet and outlet were punched and microchannel was sealed by a glass slide. The working, counter and reference electrode (WE, CE and RE, respectively) were connected to potentiostat (Volta Master 4, Hach Radiometer analytical, USA) and the sterile perfluoroalkoxy connective tubing (PFA tube 1/16, Hamilton Inc., Canada) was attached to the inlet and outlet of the device. As PDMS is known to be porous to gas molecules such as $O_2$, the microfluidic electrochemical device was placed in an anaerobic jar with 20% $CO_2$ and 80% $N_2$ (Figure S1). The tube inlet was connected to a mixer with 3 inlets from 3 different syringes with 10, 0.2, 0 mM acetate concentrations. Each of the 3 mixer inlets was connected to a one way stopcock valve to control the flow of liquid into the mixer. With flow control from syringe pumps holding gas-tight glass syringes, the nutrient concentrations were controlled based on the flow rate ratio between nutrient solutions containing different acetate concentration. All waste was contained in the anaerobic jar. To validate the ability of the setup to maintain anaerobic conditions during operation, an oxygen indicator (resazurin) was used based on the standard protocol.[1] The resazurin solution flowed into the jar, through the microfluidic device and off-chip into the waste container within the anaerobic jar. This experiment was conducted over 2 weeks without any indication of solution oxygenation. The high electrical current densities produced by the mature *Geobacter* biofilm matched the upper limits shown in the literature, thus supporting the conclusion that optimal anaerobic conditions were achieved.

Chronoamperometry and cyclic voltammetry were conducted using a commercial potentiostat (Volta Master 4, Hach Radiometer analytical, USA). All applied potentials were relative to a stable gold RE as demonstrated in previous work [2,3]. Chronoamperometry was conducted an equivalent to 0 mV vs. Ag/AgCl). All current measurements by chronoamperometry were acquired every 2.7 seconds and a 10 point smoothing algorithm was conducted to reduce the file sizes. Thus time resolution of nearly 30 s. Solution velocity through the channel was determined based on the total volumetric flow rate. Acetate concentrations delivered to the microchannel were systematically changed to observe real-time effects on current outputs via chronoamperometry.

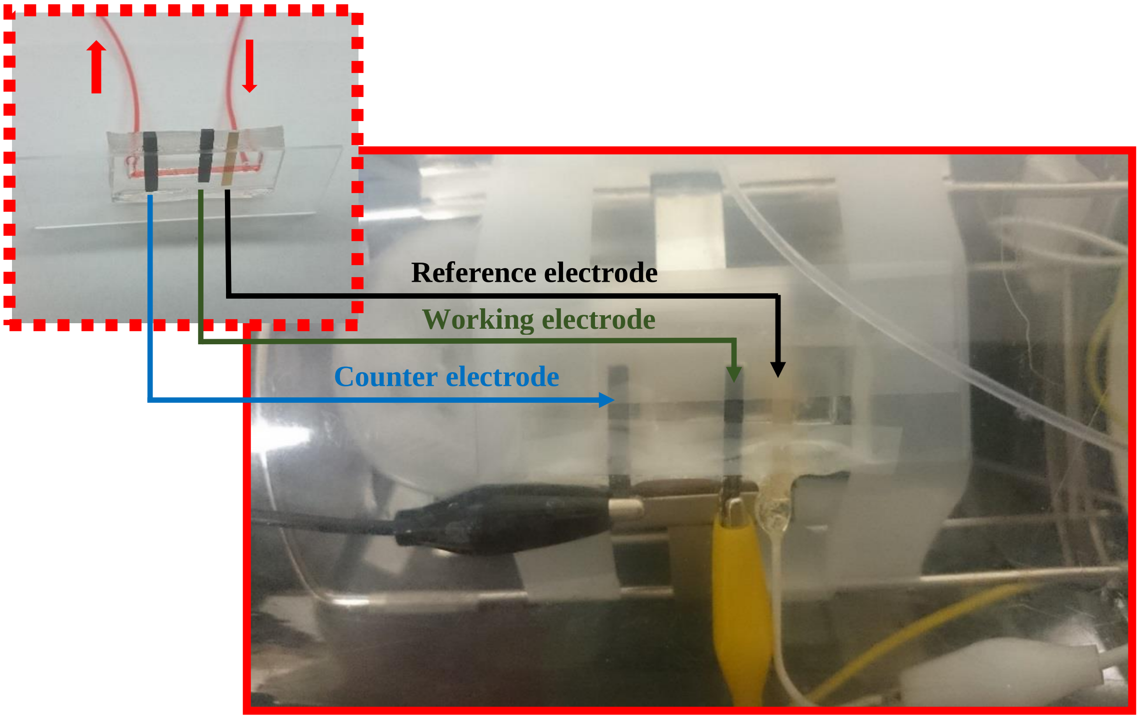


**Figure S1.** Microfluidic electrochemical cell with three-electrode setup and connections inside the anaerobic jar. Flow passes through the micro channel from inlet and touch RE, WE and CE, respectively. Inset: red solution is injected on to the device and withdrawn through connective tubing. The red liquid highlights the microfluidic channel.

### 2. Hydrodynamic considerations at and away from the working electrode

In general the flow velocity in the microchannel is calculated as

$v$ = Q/(w·h) (S1)

where h was the channel height (400 µm) and was the channel width (2 mm) and Q is the volumetric flow rate used. The growth of a biofilm on the working electrode (WE) surface will reduce the channel volume in the headspace above the electrode, which can change the local hydrodynamic properties. As discussed in the main paper the biofilm height after 300 h was assumed to be constant at 80 µm. Therefore, the channel height above the biofilm was $h_b$ = 320 µm, 80% of the clean channel height. Therefore, average velocity above the electrode-adhered biofilm, $v_b$, at a volumetric flow rate, Q (mL·h$^{-1}$) was estimated based on Eq. S1 and would have increased in proportion to the reduction free volume. Another outcome is the increase in the applied shear stress against the biofilm, compared to shear stress against the channel wall, $\tau_b$ and $\tau$ (Pa), respectively. Eq. S2a and S2b shows $\tau_b$ and $\tau$ as a function of flow rate:

$\tau_b = 6 \cdot \mu \cdot Q/(w \cdot (h-h_b)^2)$ (S2a)

$\tau = 6 \cdot \mu \cdot Q/(w \cdot (h)^2)$ (S2b)

where µ is the liquid-phase viscosity (1 × 10$^{-3}$ Pa·s) and h-$h_b$ is the headspace above the WE. Lastly, the Reynold's number was calculated both in the clean channel, Re, and above the biofilm, $Re_b$. Due to the competition between flow velocity and hydrodynamic diameter of the free volume in the clean and biofilm occupied portions of the channel, respectively, both Re and $Re_b$ were the same for a given Q. In all cases, values well-below the threshold for laminar flow.

As discussed in previous work [2,3], the gold (pseudo) RE is accurate and stable if the liquid contacting it maintains constant physiochemical conditions, most importantly the pH. To ensure this was the case, we placed the RE upstream so that any metabolites or released biofilm flows downstream. We verify that upstream proton diffusion was slower than the downstream liquid velocity ($v$) by calculating the effective diffusion velocity, $v_d$, given by Eq. S3:

$$v_d = \frac{d_d}{t} = \frac{\sqrt{2D_{H^+}t}}{t} \qquad \text{(S3)}$$

where $D_{H^+}$ (cm$^2$·s$^{-1}$) is the proton diffusion coefficient in water ($D_{H^+}$=9.31 × 10$^{-5}$ cm$^2$·s$^{-1}$). Taking t=1 s and the diffusion coefficient of a free proton as, [4] we calculated $v_d$ to be approximately 2.22 µm·s$^{-1}$ or 8.18 mm·h$^{-1}$. As this is 38 times slower than even the slowest channel flow velocity, no metabolites could reach the RE. Table S1 summarizes results for velocity, Reynold's number and shear stress above and away from the WE and for five flow rates used in the main paper as well as the proton diffusion velocity.

Table S1. Hydrodynamic parameters in the clean portions of the channel (grey) and above the biofilm (orange) for typical flow conditions used in this study.

| Q ($mL \cdot h^{-1}$) | $\bar{v}_{d^{H^+}}$ (constant) ($mm \cdot h^{-1}$) | $v$ ($mm \cdot h^{-1}$) | Re | $\tau$ (mPa) | $v_b$ ($mm \cdot h^{-1}$) | $Re_b$ | $\tau_b$ (mPa) |
|---|---|---|---|---|---|---|---|
| 0.25 | 8.2 | 312.5 | 0.065 | 1.159 | 390.6 | 0.065 | 1.811 |
| 0.5 | | 625 | 0.130 | 2.318 | 781.3 | 0.130 | 3.622 |
| 0.75 | | 937.5 | 0.195 | 3.476 | 1171.9 | 0.195 | 5.431 |
| 1 | | 1250 | 0.260 | 4.636 | 1562.5 | 0.260 | 7.243 |
| 1.5 | | 1875 | 0.390 | 6.952 | 2343.8 | 0.390 | 10.863 |

To verify that pH did not in fact change at the source, periodic measurements of the nutrient solution pH flowing into the device determined that it was constant at 7 ± 0.1 throughout the experiment. Following experiments, scanning electron microscopy (SEM) was used to verify that bacterial growth did not occur on the RE, where electrode respiration was impossible.

In summary, the RE remained sterile and the physiochemical properties of the fluid in contact with it were constant throughout the experiment, which explains its stable performance throughout the experiment, observed here and in previous work using a similar flow device [2,3].

### 3. Bacterial preparation

*G. sulfurreducens* strain PCA (ATCC 51573) was cultured in our laboratory at 30 °C using an anaerobic medium containing the following per liter of distilled water: 1.5 g $NH_4Cl$, 0.6 g $NaH_2PO_4$, 0.1 g KCl, 2.5 g $NaHCO_3$, 0.82 g $CH_3COONa$ (acetate, 10 mM), 8 g $Na_2C_4H_2O_4$ (fumarate, 40 mM), 10 mL vitamin supplements ATCC® MD-VS™, 10 mL trace mineral supplements ATCC® MD-TMS™. The filtered sterilized sodium fumarate, vitamin and trace mineral supplements were added after sterilization of all other materials at 110 °C, 20 psi pressure for 20 min at an autoclave. *G. sulfurreducens* were sub cultured in an anaerobic glove box containing the 10% $H_2$, 10% $CO_2$ and 80% $N_2$. Sub-culturing of *G. sulfurreducens* cells were done in an adjusted pH = 7 nutrient medium in the glove box and 3$^{th}$ to 8$^{th}$ subculture samples were used at electrochemical cell. Sodium fumarate and vitamins were removed from the nutrient solution for biofilm growth in electrochemical mode.

### 4. Reference electrode calibration and stability

The gold reference electrode (RE) calibration and its stability during the experiments have been deeply studied in previous research [2,3] and compared to other microfluidic three-electrode cells with different electrode setups [5]. Briefly, a 10 mM acetate nutrient medium was used to measure formal potential of cytochrome c proteins in a *G. sulfurreducens* biofilm in a three-electrode bulk electrochemical cell via CV over the course of 120 h. When the standard Ag/AgCl RE was replaced with gold RE (new or used) a shift of 410 – 396 mV in the measured potential was recorded. To verify stability of the electrochemical flow cell featuring the gold RE, a 10 mM ferricyanide in medium solution was flowed for 2 weeks. During this time no obvious formal potential change was observed. In addition, changing flow rate did not result in any changes in to the measured formal potential of ferri/ferro redox couples, even at 10 $mL\cdot h^{-1}$, which was higher than flow rates were used in biofilm measurement. The details of experiment and discussion about flow stability can be found in previous study.

### 5. Device inoculation and growth

An inoculum solution containing *G. sulfurreducens* was first introduced to the microchannel through an upstream port in the mixer unit (not shown) for 3 h at Q = 0.5 $mL\cdot h^{-1}$. The solution contained dissolved fumarate to enable extracellular electron transfer by planktonic bacteria. During inoculation, the WE was set to an oxidative electrochemical potential at 400 mV vs. Au, equivalent to 0 mV vs. Ag/AgCl, which also enabled electrode respiration for locally attached bacteria. After inoculation, the channel was exposed to a nutrient solution of [Ac] = 10 mM with no sodium fumarate, thereby ensuring the biofilm grew on the WE only. This was accomplished by opening valve $MI_1$ and setting $Q_1$ = 0.25 $mL\cdot h^{-1}$, while the other pump flow rates for were set at $Q_2 = Q_3$ = 0 $mL\cdot h^{-1}$ with valves $MI_2$ and $MI_3$ closed to prevent any back flow through the mixing unit. Under these conditions, the *G. sulfurreducens* bacteria could grow on the graphite-WE (Figure 1c) and a current was measured via chronoamperometry during the growth period (Figure S2a). As discussed in the main paper, the EAB was subjected to nutrient-limited growth media after approximately 500 h. Figure S2b, shows the reduction in current after switching from 10 mM to 0.2 mM in advance of the iterative process to find threshold concentrations. Due to the large difference in the time scale compared to Figure S2a the fluctuations appear to be lower, not due to any post-acquisition data processing (i.e. smoothing). The equilibrium current at 0.2 mM was approximately 10 µA, which was slightly above background current, proving that the EAB was still active.

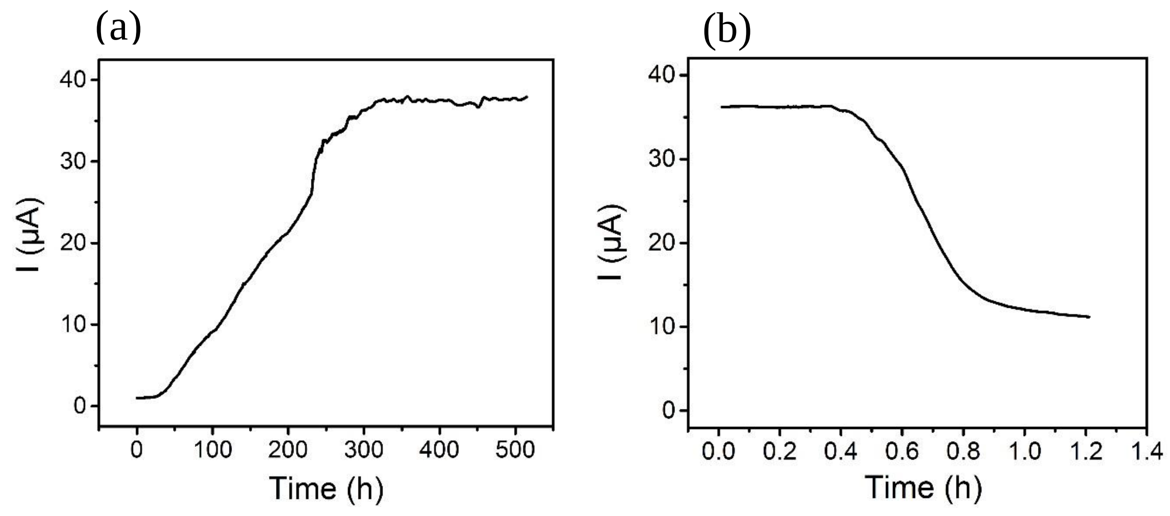

**Figure S2.** (a) *G. sulfurreducens* biofilm growth in microfluidic channel with 10 mM acetate nutrient and 0.2 mL·h$^{-1}$ flow rate for 500 h. (b) Reduction of the current by switching from 10 mM nutrient medium to 0.2 mM Ac nutrient concentration.

### 6. Identification of cytochrome c formal potential by cyclic voltammetry:

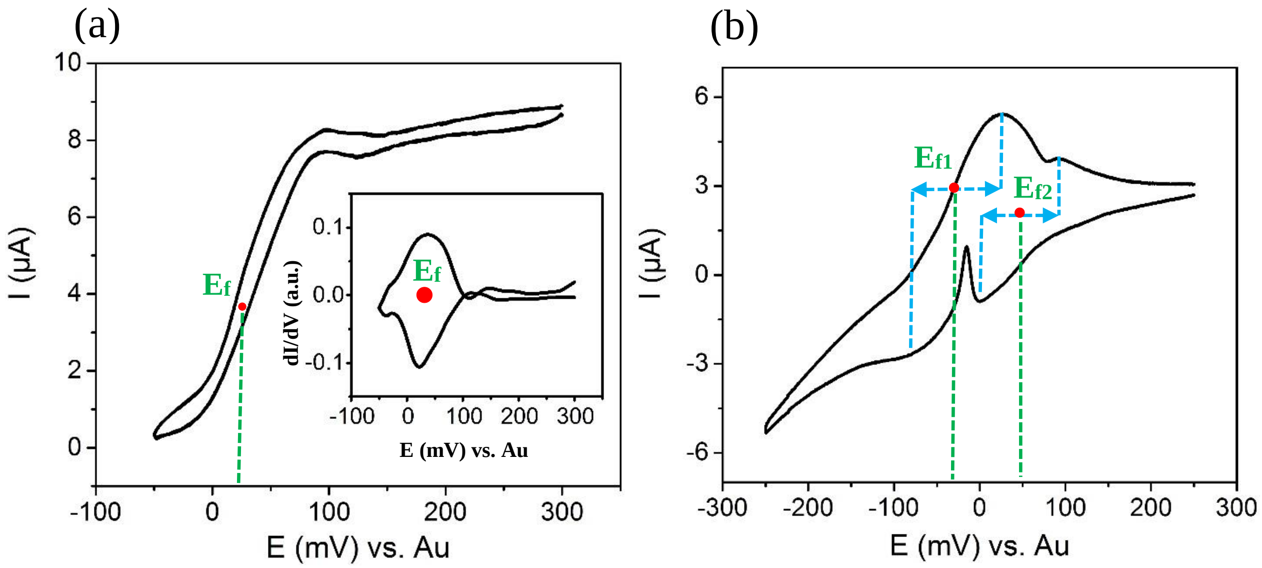


**Figure S3.** CV curve of *G. sulfurreducens* biofilm on the graphite WE at (a) turnover at 100 h and (b) non-turnover, recorded 5 hours after switching to a nutrient depleted solution. $E_f$ is formal potential of cytochromes C of *G. sulfurreducens* biofilm. Scan rates were 1 mV·s$^{-1}$ and 3 mV·s$^{-1}$ for (a) and (b), respectively.

### 7. Fluctuations in electrical current for different metabolic activity states

We demonstrate fundamental differences in the current fluctuations of an active biofilm compared to those during pseudo-activity. Figure S4a shows a typical current profile for approximately the first 200 h hours following biofilm maturation. Note the narrow range in currents shown on the y-axis. As mentioned in Section S1, the time resolution for all chronoamperometry experiments was 30 s (1/120 h$^{-1}$). This was sufficient to observe frequency components as high as 60 h$^{-1}$ according to the Nyquist sampling theorem. We used a fast Fourier transform algorithm on a typical 40 hour window (Figure S4b) compare the results to those in Figure S4c (fluctuations during pseudo-active state). A visual comparison is given via Figure S4c shows the transition to pseudo-activity after increasing acetate concentration above $[Ac]_{PA}$. The duration in both Figures S4b,c are 40 h to show the qualitative difference in oscillation periods. It is important to recognize that the y-axis scales between the two figures are very different.

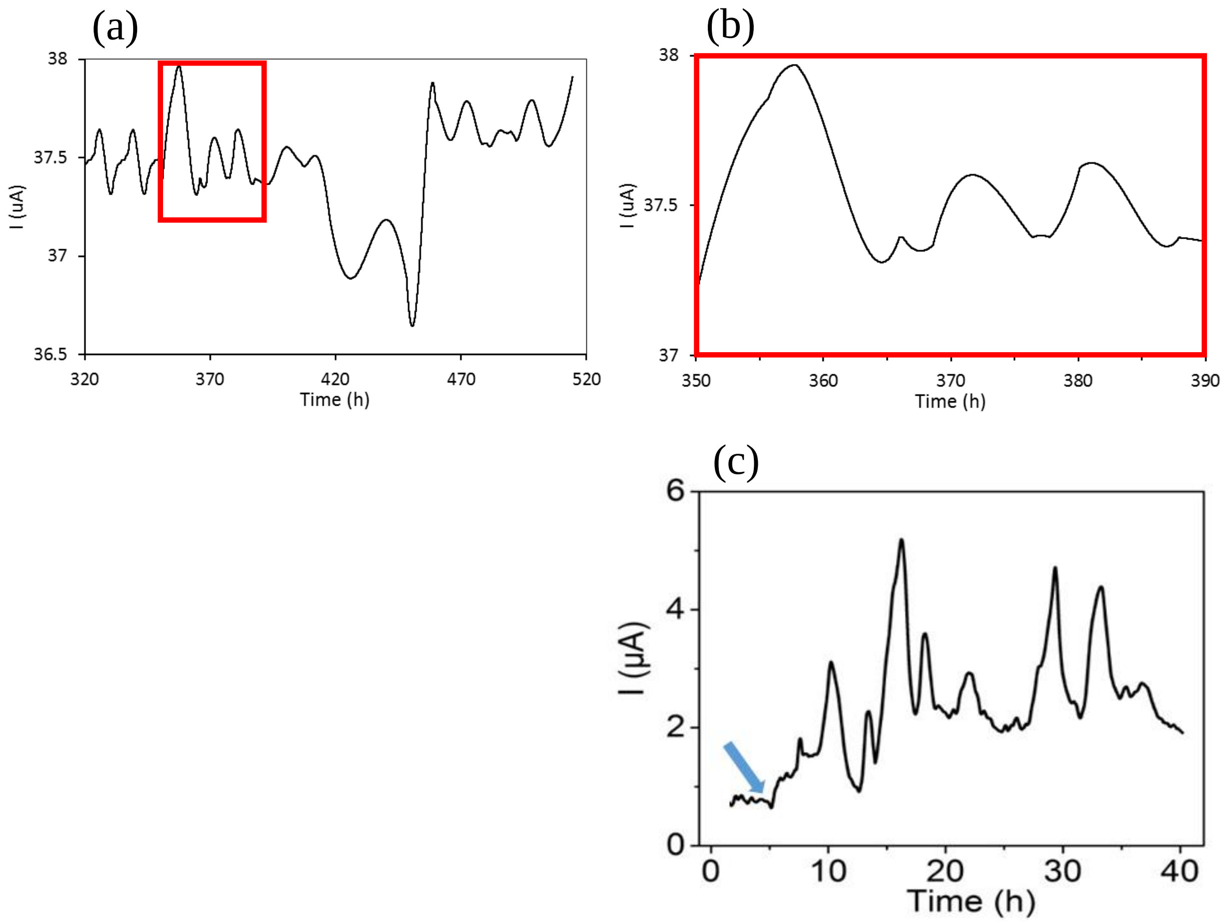


**Figure S4.** (a) CA of *G. sulfurreducens* biofilm profile for approximately the first 200 h hours following biofilm maturation. The red box shows the region of data shown in (b). (c) Nutrient solution was switched to 80 µM acetate medium (blue arrow) while the flow rate was maintained constant at 0.5 mL·h$^{-1}$. *G. sulfurreducens* biofilm shows a fluctuating (pseudo-active) state after at 80 µm acetate in 40 h exposure time.

Table S2 quantitatively tabulates key observable differences for active biofilms (exposed to 10 mM acetate, Q=0.25 mLh$^{-1}$) and pseudo-active biofilms (exposed to a variety of conditions within Zone II shown in Figure 3 in the main paper). Observables included significant upper frequency components in the current fluctuations ($f_{upper}$), average current after background current ($I_B$) subtraction ($I_{ave}$-$I_B$) and a measurement of the absolute value of peak-to-peak (P2P) current as well as the P2P changes relative to the backgrounded adjusted average current. The results demonstrate very stark differences which can be summarized in two general observations: (1) current fluctuations from pseudo-active biofilms have high frequency components are that are approximately 40 times faster than those for active ones and (2) the relative intensity of the current fluctuations versus the average current output for pseudo-active is about 50 times higher than for active biofilms. Together these observations point to a fundamental difference in behaviour of biofilms in a pseudo-active state compared to the active state.

Table S2. Comparison of current fluctuations between active and pseudo-active biofilms.

| | | | **P2P fluctuations** | |
|---|---|---|---|---|
| **Biofilm state** | **$f_{upper}$ ($h^{-1}$)** | **$I_{ave}$-$I_B$** | **Raw (µA)** | **Relative (%)** |
| **Active** | 0.05 | 36.5 | 1.5 | 4.1 |
| **Pseudo active** | 2 | 2.6 | 5.4 | 207.7 |

### 8. Efficiency of acetate digestion

As noted in the main paper, when supplied concentrations at the activity threshold $[Ac]_A$.the final acetate concentration $[Ac]_f$ in the range of 10 to 20 µM indicated efficient oxidisation. This was notable because the supply concentrations were lower than in usable concentrations in bulk systems and as well because liquid/biofilm contact times were short, ranging from just 6 to 35 seconds, depending on the flow rate. Based on Equation 4 in the main paper (reproduced here as Equation S5) acetate conversion efficiency ($\varepsilon_{Ac}$) from initial concentration, $[Ac]_0$, to final concentration, $[Ac]_f$ is calculated:

$$\varepsilon_{Ac} = ([Ac]_0-[Ac]_f)/[Ac]_0$$

Using data in the main Figure 3 for the different $[Ac]_A$, $[Ac]_f$ pairs for each Q value we obtained conversion efficiencies shown in Figure S5a. The same procedure was used to generate $\varepsilon_{Ac}$ for $[Ac]_{PA}$ and for two higher concentrations, [Ac]=0.3 and 10 mM. It should be restated that the absolute concentration for the two thresholds $[Ac]_A$ and $[Ac]_{PA}$ changed with Q, whereas the tests at higher concentrations were held constant during changes to Q. Plotting $\varepsilon_{Ac}$ versus Q in Figure S5a shows that acetate conversion is most efficient at activity threshold concentrations. This is expected for high concentration solutions above the traditional zero order concentration regeims because current does not increase in proportion to concentration and more time is required to reduce concentrations. However, even at the relatively low concentration of 0.3 mM, $\varepsilon_{Ac}$ is notably worse than at either of the threshold concentrations. The general reduction in $\varepsilon_{Ac}$ with increasing Q is expected since contact time reduced. We note that the time normalized efficiency actually increases with flow rate (Figure S5b). This is likely due to flow effects that both reduce the diffusion barrier around the biofilm and enable deeper penetration of the nutrient flow into the biofilm, effectively increasing accessibility of nutrients to bacteria in lower biofilm strata.

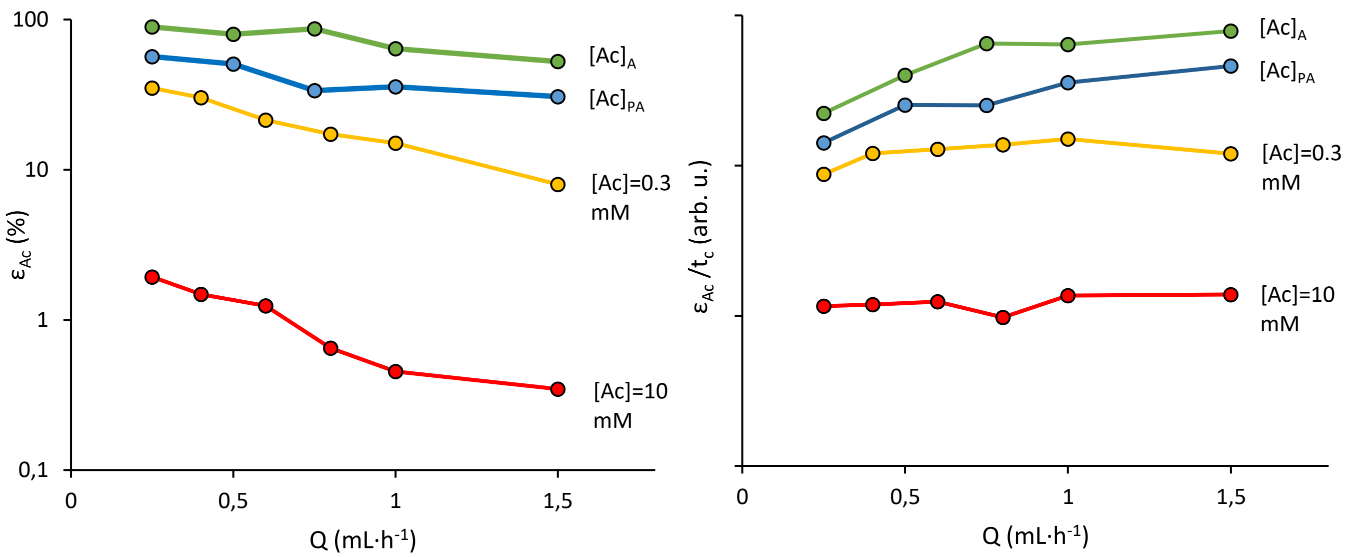


**Figure S5**. (a) Acetate conversion efficieny ($\varepsilon_{Ac}$) and (b) acetate conversion efficiency normalized by contact time ($\varepsilon_{Ac}/t_c$) as a function of Q for concentrations indicated. Exact values of $[Ac]_A$ and $[Ac]_{PA}$ are given in Figure 3 in the main paper. Both y-axes are plotted on a log-scale.
